# CellFuse Enables Multi-modal Integration of Single-cell and Spatial Proteomics data

**DOI:** 10.1101/2025.07.23.665976

**Authors:** Abhishek Koladiya, Zinaida Good, Sricharan Reddy Varra, Sean C Bendall, Kara L. Davis

**Affiliations:** Division of Hematology, Oncology, and Stem Cell Transplant and Regenerative Medicine, Department of Pediatrics, Stanford University, CA; Division of Immunology and Rheumatology, Department of Medicine, Stanford University; Center for Biomedical Informatics Research, Department of Medicine, Stanford University; Parker Institute for Cancer Immunotherapy at Stanford, Stanford University; Department of Pathology, Stanford University, Stanford, CA; Center for Cancer Cell Therapy, Stanford Cancer Institute, Stanford University School of Medicine, Stanford, CA

## Abstract

Single-cell and spatial proteomic technologies capture complementary biological information, yet no single platform can measure all modalities within the same cell. Most existing integration methods are optimized for transcriptomic data and rely on a large set of shared, strongly linked features, an assumption that often fails for low-dimensional proteomic modalities. We present CellFuse, a deep learning-based, modality-agnostic integration framework designed specifically for settings with limited feature overlap. CellFuse leverages supervised contrastive learning to learn a shared embedding space, enabling accurate cell type prediction and seamless integration across modalities and experimental conditions. Across a range of datasets including healthy PBMCs, bone marrow, CAR-T–treated lymphoma, and healthy and tumor tissues—CellFuse consistently outperforms existing methods in both integration quality and runtime efficiency. It maintains high accuracy even in the presence of missing markers and rare cell types, and performs robustly in cross-dataset comparisons, making it a powerful tool for scalable and high-fidelity single-cell data integration in basic and translational research.

## Main

The rapid advancement of single-cell technologies has revolutionized our ability to study complex biological systems at unprecedented resolution^1^. Today, it is possible to individually measure genomic, epigenomic, transcriptomic, and proteomic states in single cells, ushering in a new era of systems-level biology. Single-cell RNA sequencing (scRNA-seq) and targeted proteomics have become essential for characterizing cellular identity and function. While scRNA-seq can capture thousands of gene transcripts per cell, it often suffers from data sparsity, particularly for low-abundance genes. In contrast, single-cell proteomics captures proteins-more stable and functionally relevant indicators of cell state-that influence cellular behavior through regulation at multiple levels, including epigenetic, transcriptional, and translational pathways. The capability to characterize proteomic states at single-cell resolution, both in dissociated cells and intact tissue contexts, represents a significant leap forward in biology. Suspension-based single-cell proteomic modalities, such as flow, spectral, and mass cytometry (CyTOF), have progressed from measuring a handful of markers^2^ to routinely detecting over 50 proteins^3, 4^. Emerging sequencing-based modalities such as antibody sequencing (AbSeq)^5^ and cellular indexing of transcriptomes and epitopes by sequencing (CITE-Seq)^6^ have further expanded this capacity, enabling the simultaneous quantification of more than 200 proteins alongside transcriptomic data. Meanwhile, spatial proteomic platforms—including co-detection by indexing (CODEX)^7^, multiplexed ion beam imaging by time of flight (MIBI-TOF)^8^, and imaging mass cytometry (IMC)^9^—now allow for the high-plex measurement of more than 50 proteins directly within tissue sections, preserving spatial context and enabling deeper insights into the organization and function of complex cellular environments.

As the proteomic features measured across modalities overlaps, tools that can obtain cell type annotations from well-curated reference modalities (e.g. CyTOF or CITE-Seq) to annotate cells from query modalities (e.g. CODEX) offer a valuable opportunity for cross-modal integration— leveraging extensive cellular phenotyping from dissociated single-cell methods to inform the spatially resolved data and vice versa. However, a major hurdle in this process lies in aligning cells across modalities, especially given that most existing integration algorithms are optimized for transcriptomic data, which typically include thousands of shared features^10–12^. In contrast, single-cell proteomic datasets contain far fewer overlapping features, rendering conventional approaches such as mutual nearest neighbors (MNN) and canonical correlation analysis (CCA) ineffective. These methods often struggle in low-dimensional spaces, failing to capture the full phenotypic diversity of the data, which leads to poor cell matching, distorted embeddings, and suboptimal integration. This underscores the need for specialized approaches tailored to the unique challenges of integrating single-cell and spatial proteomic data.

To overcome these challenges, we developed CellFuse, a deep learning–based method designed for integrating single-cell proteomics data. Given an annotated suspension or spatial proteomic dataset with labeled cell types for reference, CellFuse predicts cell types in a query dataset based on shared protein measurements and performs data integration. We systematically benchmarked CellFuse across single-cell proteomic modalities, including CyTOF and CITE-seq data derived from bone marrow (BM) and peripheral blood (PB) samples. In our evaluations, CellFuse consistently outperformed existing integration methods in both cell type preservation and batch correction, demonstrating superior accuracy in maintaining biologically meaningful structures while minimizing technical variation across datasets. We next demonstrated the analytical capabilities enabled by CellFuse through three distinct applications: (1) multi-modal integration of CAR T cell datasets to identify disease-associated cell populations, (2) annotation of healthy colon tissue profiled by CODEX from the Human BioMolecular Atlas Program (HuBMAP) and breast carcinoma samples profiled by IMC, and (3) integration of single-cell CyTOF data with spatial MIBI-TOF data to resolve the cellular architecture of the tumor microenvironment in colon carcinoma. To ensure broad accessibility, we implemented the CellFuse algorithm as an open-source R package available at: https://github.com/karadavis-lab/CellFuse

## Results

### Overview of CellFuse

The initial input for CellFuse is a protein expression matrix derived from either single-cell suspension or spatial proteomic data that utilizes antibodies to label cells. For convenience, we refer to this input as the reference data. The reference data comprises two components: (1) a cell-by-protein matrix that captures the measured abundance of each protein across all cells, and (2) a corresponding set of ground truth annotations that specify the cell type of each cell in the dataset. CellFuse operates in a three-stage framework that leverages the reference data to predict cell type labels for unannotated query data and subsequently integrate it with the reference data (Fig. 1).

**Fig. 1:**
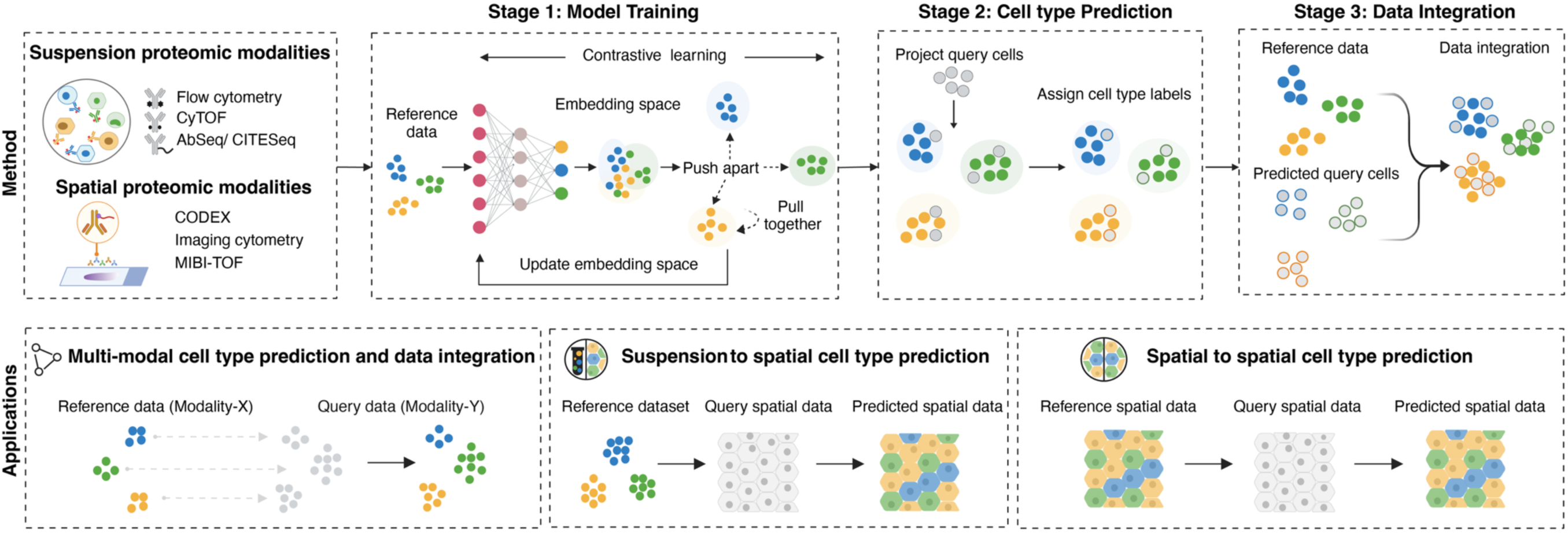
Schematic of the CellFuse analysis pipeline. Single-cell proteomic profiles can be generated using suspension-based technologies—such as flow cytometry, CyTOF, AbSeq, and CITE-seq—or spatial platforms including CODEX, imaging mass cytometry, and MIBI-TOF. Given a reference dataset from either modality, CellFuse begins with Stage 1 (Model Training), where a neural network is trained using contrastive learning to construct an embedding space that places phenotypically similar cell types closer together and dissimilar types farther apart. This embedding is iteratively refined to enhance biological separability. In Stage 2 (Cell Type Prediction), query cells—sourced from either suspension or spatial proteomic data—are projected into the learned space, and cell type labels are assigned based on proximity to reference cells. Finally, Stage 3 (Data Integration) integrates the predicted query cells with the reference dataset to enable downstream joint analyses. CellFuse has three major applications: (1) multi-modal cell type prediction and integration, enabling reference data from CyTOF to annotate and integrate query data from CITE-seq; (2) cross-platform cell type transfer from suspension-based to spatial proteomic data, such as from CyTOF to CODEX or MIBI-TOF; and (3) spatial-to-spatial annotation transfer, including MIBI-TOF to MIBI-TOF. CyTOF, mass cytometry; AbSeq, Antibody Sequencing; CITESeq, cellular indexing of transcriptomes and epitopes by sequencing; CODEX, CoDEtection by indeXing; MIBI-TOF, Multiplexed Ion Beam Imaging by Time Of Flight (MIBI-TOF)

In the first stage, a neural network is trained on the reference data using a supervised contrastive learning framework. This training strategy encourages the model to learn an embedding space in which cells of the same type are projected closer together, while cells of different types are mapped further apart. The contrastive loss function leverages the ground truth annotations to define positive (same-type) and negative (different-type) cell pairs, enabling the model to capture biologically meaningful relationships across protein expression profiles. The resulting encoder produces a low-dimensional representation that preserves cell identity while reducing noise and redundancy in the high-dimensional input space. To improve generalization and prevent overfitting, 40% of the reference data is held out as a validation set during training, which provides a robust balance between model optimization and evaluation stability. Training proceeds for a user-defined number of epochs or until early stopping criteria are met based on the stability of the validation loss. Upon convergence, the trained encoder is saved and used for downstream prediction.

In the second stage, the trained encoder is used to project unannotated query cells into the same low-dimensional embedding space learned from the reference data. Once embedded, each query cell is assigned a predicted label based on its proximity to the labeled reference cells using a nearest-neighbor classification approach. For each query cell, the most frequent label among its nearest neighbors is selected as the predicted cell type, and a confidence score is computed based on the agreement among neighbors. This procedure enables robust and scalable annotation of query cells.

In the third stage, following label prediction and initial embedding, a distributional integration step is performed to further align the query data with the reference. Although the query cells are embedded in the same low-dimensional space as the reference, technical or modality-specific differences may still result in mismatches in their distributional profiles. To address this, both reference and query datasets are projected into a shared principal component (PC) space, using the PCA basis computed from the reference data. Within this space, the distribution of the query projections is adjusted to better align with the reference along each principal component. The integrated query profiles are then projected back into the original protein feature space using the inverse PCA transformation. This step enhances the consistency between query and reference data at the distributional level, improving downstream interpretability and enabling more accurate joint visualization and analysis.

### Benchmarking and integration of multimodal suspension datasets

To evaluate the performance of CellFuse, we utilized suspension proteomic datasets from two biological sources: BM and PBMC (Fig. 2A). The BM dataset includes cells profiled using CITE-seq with a 25-marker antibody panel^10^ and CyTOF with a 32-marker panel^13^, with 12 markers and 8 cell types shared across modalities. The PBMC dataset comprises cells analyzed using a 45-marker CyTOF panel^14^ and a 228-marker CITE-seq panel^15^, with 27 overlapping markers and 12 common cell types between the two modalities. We benchmarked the performance of CellFuse against existing single-cell integration methods, including Seurat^10^, FastMNN^12^, and Harmony^11^. To ensure a fair evaluation of generalizability, we benchmarked both directions of integration: using CyTOF as the reference and CITE-Seq as the query for the BM dataset, and vice versa for the PBMC dataset, where CITE-Seq was used as the reference and CyTOF as the query.

**Figure 2:**
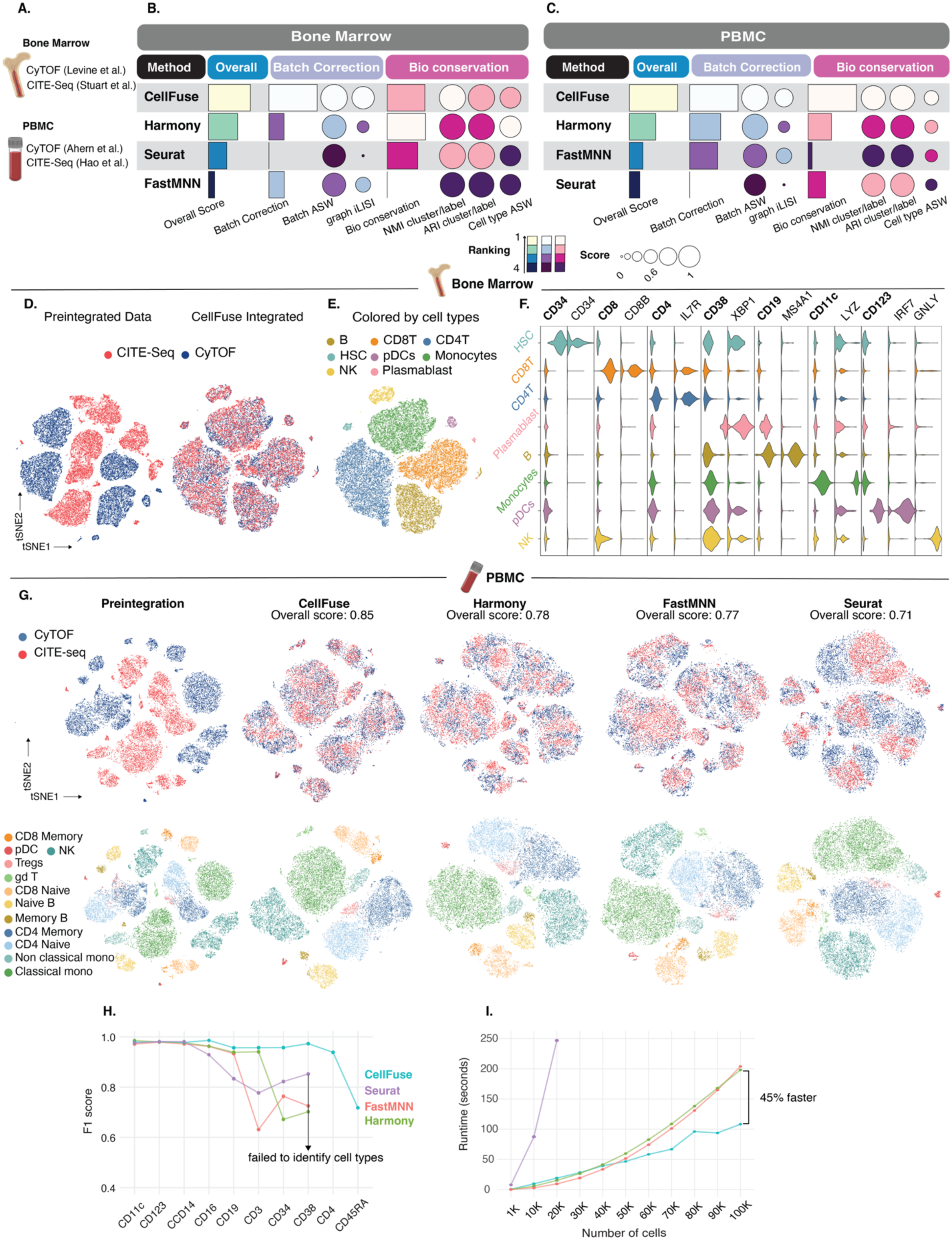
Benchmarking of CellFuse and other integration methods on ground-truth BM and PBMC data. **A.** Two different single-cell datasets, generated by different technologies, were benchmarked. **B. C.** Benchmarking of CellFuse’s performance against Harmony, Seurat and FastMNN using BM (B) and PBMC (C) dataset. Metrics are divided into batch correction (violet) and bio-conservation (pink) categories. Overall scores (blue) are computed using a 40/60 weighted mean of these category scores, suggested by Luecken et al (Nature Methods 2021). **D.** t-SNE plots of individual cells from BM dataset colored by modality, preintegrated (left) and CellFuse integrated (right). **E.** t-SNE plots of CellFuse integrated BM dataset colored by cell type. **F.** Violin plot showing protein (bold) and gene expression per cell type post CellFuse integration. **G.** t-SNE plots visualizing the integration of the PBMC dataset. From left to right: pre-integration, CellFuse, FastMNN, Seurat, and Harmony. The top row is colored by modality, highlighting the degree of batch mixing, while the bottom row is colored by cell types. **H.** Line plot showing F1 scores for each integration method (CellFuse, Seurat, FastMNN, Harmony) as overlapping protein features are sequentially dropped from the Bone Marrow dataset. **I.** Runtime comparison of integration methods across increasing subsampled cells from the PBMC dataset. BM, bone marrow; PBMC, peripheral blood mononuclear cells; B, B cells; HSC, H, hematopoietic stem cells; NK, natural killer cells; pDCs, plasmacytoid dendritic cells; CD4 Memory, CD4 Memory T cells; CD4 Naive, CD4 Naive T cells; Treg, T regulatory cells; CD8 Memory, CD8 Memory T cells; CD8 Naive, CD8 Naive T cells; gdT, gamma delta T cells; Memory; Memory B cells; Naive B, Naïve B cells; classical mono, classical monocytes; Non classical mono, Non classical monocytes.

For quantitative evaluation of integration performance, we used the single-cell integration benchmarking (scIB) tool^16^. We selected graph inverse Local Inverse Simpson’s Index (iLISI) and batch Average Silhouette Width (ASW) metrics to assess batch correction and Normalized Mutual Information (NMI), Adjusted Rand Index (ARI) and cell type ASW metrics to evaluate biological conservation.

Across both BM and PBMC datasets, CellFuse achieved the highest overall score, indicating robust correction of batch effects and preservation of biologically meaningful structure (Fig. 2B, C and Supplementary Fig. 1). In the BM dataset, CellFuse outperformed other methods in both batch correction (high Batch ASW and iLISI) and biological conservation (notably high NMI and ARI scores). Harmony and Seurat performed comparably in biological metrics but showed reduced effectiveness in removing batch effects. FastMNN had consistently lower performance across all categories. In the PBMC dataset, CellFuse again ranked first overall, achieving near-perfect batch ASW and top scores in cell type conservation (Fig. 2C and Supplementary Fig. 1B). Harmony showed improved batch correction relative to its performance in BM but lagged in cell type ASW and cluster separation. Seurat continued to excel in biological conservation (NMI and ARI), but remained weak in addressing batch effects. FastMNN exhibited moderate batch correction but had low scores for biological conservation, particularly NMI and ARI.

These results can be noted by visual inspection of BM data pre-integration (Fig. 2D, left) and post CellFuse integration (Fig. 2D, right and 2E) using t-distributed stochastic neighbor embedding (t-SNE). CellFuse effectively aligned the two modalities, achieving well-mixed batches and distinct separation of cell types. In contrast, comparison methods showed poor modality mixing (Supplementary Fig. 2), resulting in suboptimal batch correction compared to CellFuse. Given CellFuse’s superior performance in integrating CITE-Seq (query) with CyTOF (reference) using only protein features, we next examined whether the transcriptomic profiles of the integrated cells aligned with the expected protein expression patterns. We observed strong concordance between expected protein and RNA transcript expression measured by CITE-Seq for each cell type (Fig. 2F and Supplementary Fig. 2). This highlights the utility of CellFuse in scenarios where only the cytometry reference is labeled, enabling reliable annotation and integration of CITE-Seq query data based solely on shared protein features.

We observed similar strong performance of CellFuse in integration of the PBMC dataset (Fig. 2G and Supplementary Fig. 3). Prior to integration, cells from the two modalities formed distinct, modality-specific clusters with poor overlap. Following integration, CellFuse achieved near-complete mixing of the two modalities while preserving clear separation between canonical immune cell types. In contrast, Harmony, FastMNN, and Seurat showed varying degrees of incomplete mixing and distortion of cell-type boundaries. For example, γδT cells and Tregs exhibited fragmented or ambiguous structure. The CellFuse integration not only achieved the highest overall score (0.85) but also maintained both batch correction and biological conservation, making it a robust method for integrating PBMC data across modalities using shared protein features. Together, these results highlight CellFuse as a robust and generalizable integration strategy across biological sources.

In addition to the scIB benchmarking results, we further evaluated CellFuse’s integration performance through two complementary analyses: (1) sequential feature dropping to assess robustness to missing markers, and (2) runtime profiling to evaluate computational efficiency across datasets. For the sequential feature dropping analysis, we used the BM dataset, which contains 12 shared markers between CyTOF and CITE-Seq and removed 10 of these markers in a fixed order and evaluated performance at each step. As shown in Fig. 2H, both FastMNN and Seurat experience a sharp decline in F1 score after the removal of five overlapping features, ultimately failing to identify cell types beyond the exclusion of CD38. While Harmony demonstrates slightly better resilience, its performance also steadily deteriorates with continued feature loss. Notably, CellFuse demonstrates remarkable robustness, maintaining stable F1 scores when only two out of twelve shared markers remain. To evaluate computational scalability, we used the PBMC dataset, which contained a large number of cells from both CyTOF (n = 413,019) and CITE-seq (n = 130,278). We subsampled increasing numbers of cells per modality (from 1,000 to 100,000) and recorded the runtime of each integration method. CellFuse consistently outperformed all other approaches, demonstrating linear scalability with dataset size and completing integration of 100,000 cells in just 108.21 seconds—approximately 45% faster than the next fastest method (Fig. 2I).

### Cross-modality integration of CAR T cell data using CellFuse recovers clinically relevant cells

Robust integration of multi-modal datasets enables validation across studies to evaluate clinically relevant populations. For example, integrating data from CyTOF and CITE-Seq allows researchers to leverage complementary information enabling identification of disease-associated cell states with both proteomic and transcriptomic resolution. To demonstrate CellFuse’s ability to perform clinically meaningful integration, we analyzed a CAR T cell dataset from Good et al.¹⁷, in which PBMCs were collected from large B cell lymphoma (LBCL) patients treated with axicabtagene ciloleucel (axi-cel), a CD19-targeted CAR T cell therapy.

In this study, PBMCs were profiled using both CyTOF and CITE-seq at two time points: pre-infusion and seven days post-infusion. The CyTOF analysis identified a subset of CD4⁺Helios⁺CD57⁻ CAR T cells enriched in patients with progressive disease. Further CITE-seq analysis characterized these cells as Treg-like based on surface protein markers (CD4, CD25, CD28) and transcripts (CD4, FOXP3, IL2RA). However, cross-modality identification of this clinically relevant population poses two key challenges: (1) the post-infusion Day 7 CyTOF data exhibited strong batch effects (Fig. 3C and Supplementary Fig. 4A), and (2) Helios, a key intracellular Treg marker, was absent in the CITE-seq dataset, limiting direct cross-platform annotation. To overcome these challenges, we used pre-infusion CyTOF data—where batch effects were minimal and the Helios marker was available—as a reference, and applied CellFuse to integrate it with day 7 CyTOF and CITE-seq datasets (Fig. 3A, B). CellFuse effectively mitigated batch effects while preserving biologically meaningful structure, resulting in well-separated cell populations post-integration (Fig. 3D and Supplementary Fig. 4B–D). Compared to the original study-defined Treg-like population (meta-cluster 4), CellFuse-predicted Treg frequencies were significantly correlated in both progressive disease (PD) (Spearman R = 0.90, p < 2.2e−16) and complete response (CR) (Spearman R = 0.63, p = 0.04) groups (Fig. 3E), validating that CellFuse recapitulates clinically relevant cell distributions^17^.

**Figure 3:**
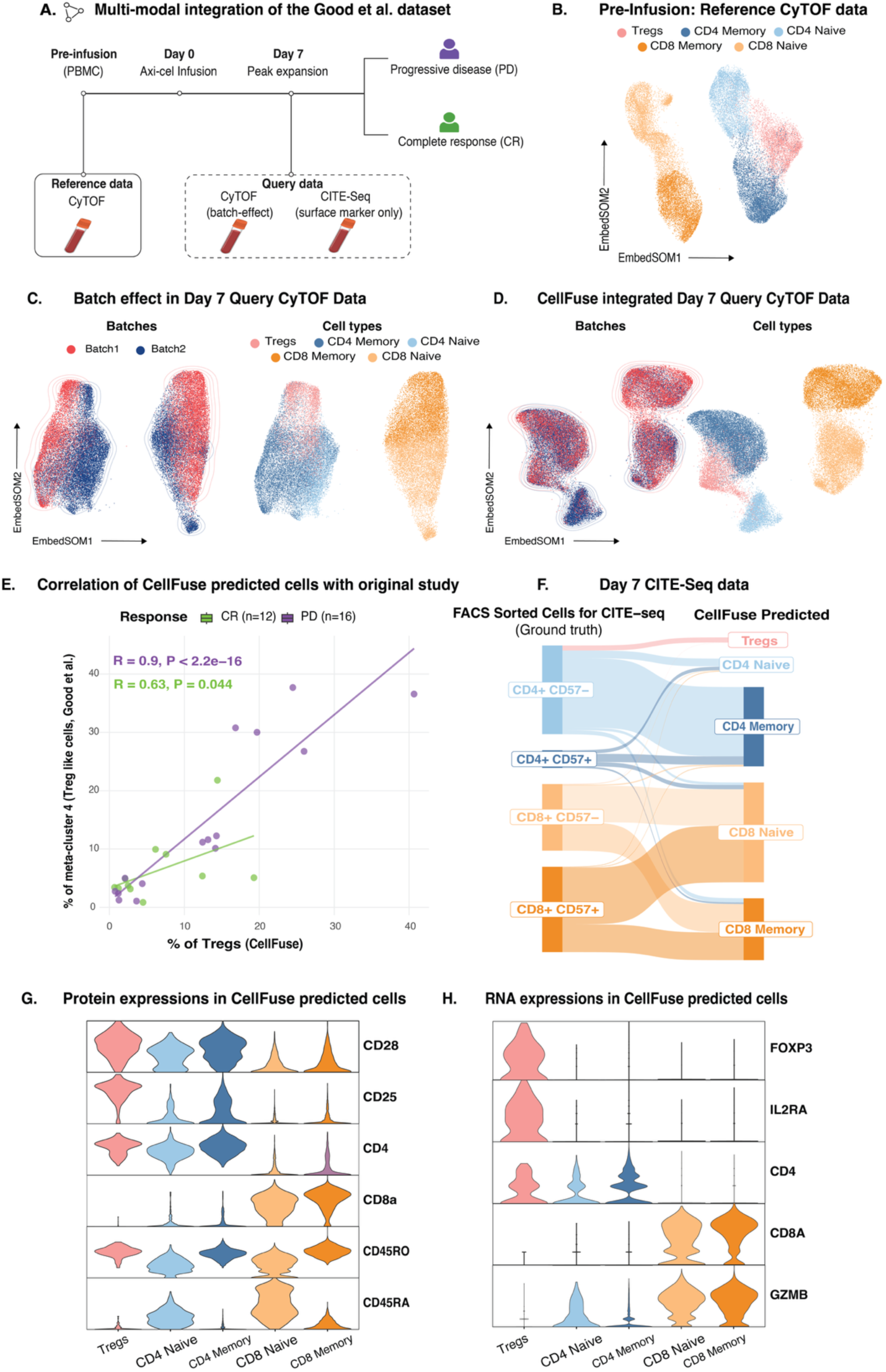
Multi-modal integration of CAR T cell data using CellFuse reveals disease-associated cell types. **A.** Schematic representation of Axi-Cel treated LBCL patients and CellFuse analysis. CellFuse model was trained on T cells from pre-infusion PBMCs to predict and integrate CART cells at day 7 post-infusion measure by CyTOF and CITE-seq. **B.** EmbedSOM plot of Day 7 CART cells displaying batch effects, colored by batch. **C.** EmbedSOM plot of pre-integrated CART cells predicted by CellFuse. **D.** EmbedSOM plot of CellFuse integrated CART cells colored by batch (left) and cell-type (right) **E.** Correlation plot of the frequency of Tregs-like cells predicted by CellFuse (x-axis) and Good et al. (y-axis) in CR or PD patients at Day 7. **F.** Sankey diagram illustrating the predictions of FACS-sorted cells measured by CITE-seq (center), with CellFuse (left) using protein features and Seurat (right) using both protein and RNA features. **G. H.** Violin plots showing selected surface protein and gene expressions of CellFuse predicted cells. Box plots in F show quartiles with a band at median; whiskers indicate 1.5× interquartile range; and all observations are overlaid as dots. PBMC, peripheral blood mononuclear cells; CD4 Memory, CD4 Memory T cells; CD4 Naive, CD4 Naive T cells; Treg, T regulatory cells; CD8 Memory, CD8 Memory T cells; CD8 Naive, CD8 Naive T cells.

Next, we focused on identifying Tregs in the CITE-Seq data using pre-infusion CyTOF data as a reference. However, since Helios protein was absent in the CITE-Seq data, we relied solely on surface markers for Treg identification. Notably, CITE-Seq cells had been previously FACS-sorted into CD4/CD8/CD57-based populations^17^. CellFuse’s predicted annotations showed strong concordance with the experimental labels (Fig. 3F), with the Tregs aligning specifically with CD4⁺CD57⁻ sorted cells. Furthermore, CellFuse-identified Tregs expressed canonical Treg proteins (CD4, CD25, CD28) and transcripts (CD4, FOXP3, IL2RA) (Fig. 3G–H), mirroring the immunophenotype reported in the original study.

Together, these results underscore CellFuse’s ability to perform robust integration across platforms and batches, enabling reliable identification of clinically important immune subsets even when key markers are missing in one modality. These findings highlight CellFuse’s utility in translational research settings, where robust batch correction and accurate cross-platform annotation are essential for identifying clinically relevant cell populations and linking them to therapeutic outcomes.

### CellFuse accurately annotated spatial proteomics data across tissue and modalities

Robust and scalable cell type annotation is critical for spatial proteomics, particularly in large cohorts where consistent identification of fine-grained cell types across donors is essential for comparative and clinical analyses. To assess CellFuse’s ability to address this challenge, we applied it to two complex spatial datasets: CODEX imaging of normal human intestine and IMC imaging of human breast cancer samples^18, 19^. The CODEX dataset, generated by the HuBMAP consortium, includes samples from two donors—one with expert-annotated cell types and one unannotated—spanning eight distinct intestinal regions (four from the small bowel (SB) and four from the colon (CL); Fig. 4A). We used cells from one SB and one CL region from the annotated donor as reference and predicted cell types in the remaining three regions of each tissue. To benchmark performance, we compared CellFuse to two established spatial cell phenotyping methods, CELESTA^20^ and Astir^21^, as well as two classification methods, support vector machine (SVM)^22^ and Seurat^10^, using the same reference data for training across all methods to ensure a fair comparison. CellFuse outperformed all comparison methods, achieving the highest median F1 score (0.81) while also being the most computationally efficient, completing predictions in just 80 seconds (Fig. 4B). Importantly, CellFuse showed superior accuracy for rare cell populations, such as CD57⁺ enterocytes (n = 78), Paneth cells (n = 177), and Neuroendocrine cells (n = 1,538), which were not detected by existing cell phenotyping methods and classification methods failed to achieve an F1 score above 0.5 for these populations (Supplementary Fig. 6A). CELESTA showed the lowest F1 score and annotated most cells as “Unknown,” a similar observation previously reported by Huynh et al. ^23^. To further validate these findings, we visually compared CellFuse predictions to expert annotations in one colonic region. The predicted and ground truth cell-type distributions showed strong spatial concordance, highlighting CellFuse’s ability to recover biologically meaningful cell identities in situ (Fig. 4C).

**Figure 4:**
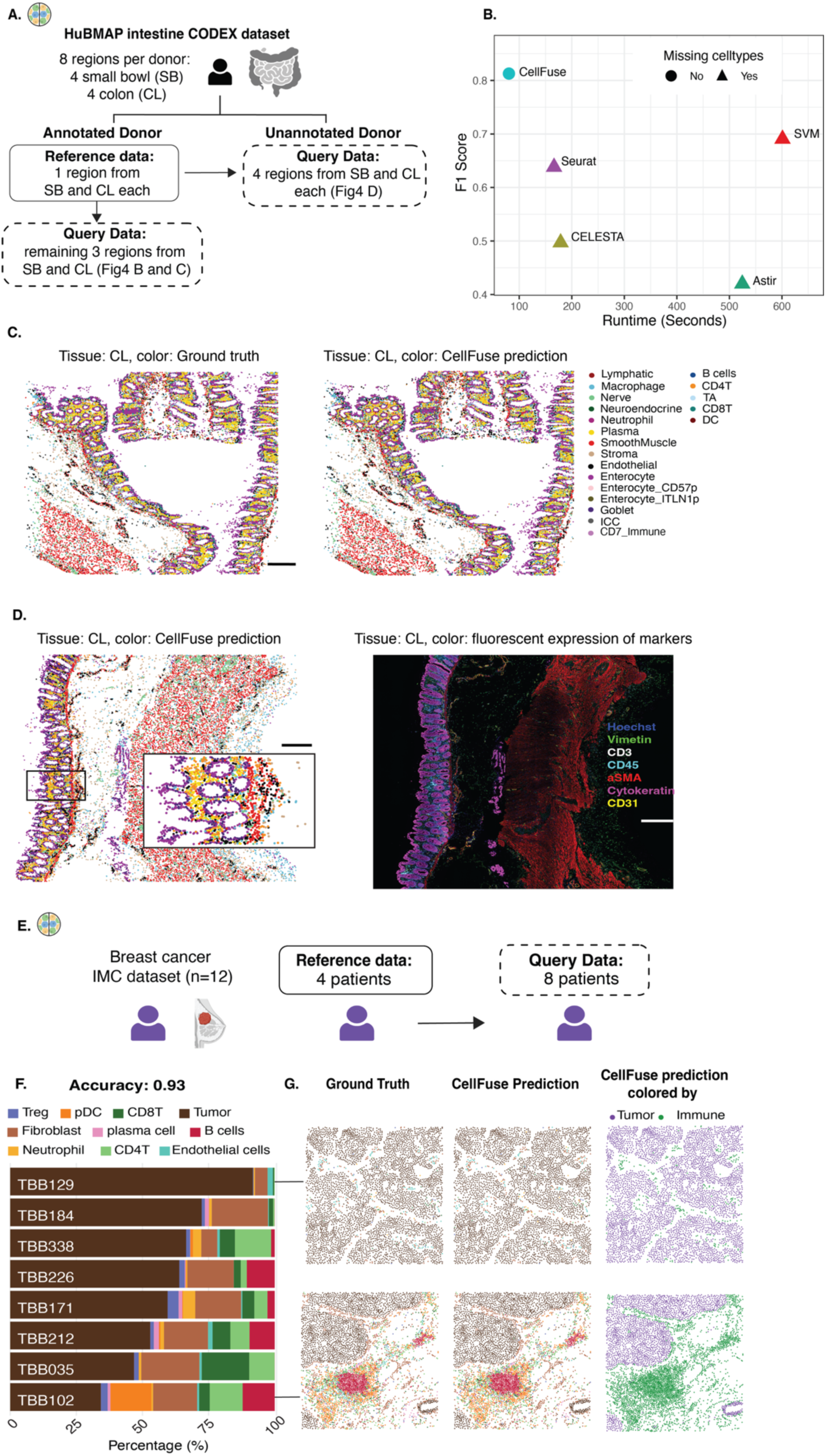
CellFuse enables cell type prediction across tissue regions and donors within HuBMAP CODEX and IMC breast cancer data. **A.** Schematic representation of CODEX human intestine dataset analysis by CellFuse. The dataset consists of one annotated (blue) and one unannotated (red) donor, each with four intestine regions. CellFuse model was train on one region from annotated donor. **B.** Benchmarking CellFuse against alternative approaches based on F1 score and runtime. **C.** Comparison of ground-truth (left) and CellFuse predicted (right) cell type. Scale bar: 100 μm **D.** Left: CellFuse-predicted cell type annotations for an independent donor, visualized on the spatial map. Scale bar: 100 μm. Right: Corresponding fluorescence image from the same donor showing protein marker staining. Channels are color-coded: Hoechst (blue), Vimentin (green), CD3 (white), αSMA (red), CD45 (cyan), Cytokeratin (magenta), and CD31 (yellow). Scale bar for the overview image: 250 μm. **E.** Illustrative overview of IMC breast cancer data analysis. CellFuse model was trained on cells from 4 patients, and predictions were made for the remaining 8 patients. **F.** Barplot showing frequency of CellFuse predicted cell types from breast cancer IMC dataset **G.** Cells from patient TBB129 (top) and TBB102 (bottom). In both cases left plots indicates ground truth cells, middle plot CellFuse predicted cells and the plots in right are CellFuse predicted cells categorized in tumor and immune cell types. Scale bar: 200 μm CODEX, CO-Detection by indexing; SVM, Support Vector Machine; CD4T, CD4 T cells; CD8T, CD8 T cells, DC, dendritic cell; ICC, interstitial cells of Cajal; TA, transit amplifying cell; pDC, plasmacytoid DCs; Tregs, regulatory T cells; IMC, Imaging Mass Cytometry.

Next, we evaluated the robustness of CellFuse across donors, again utilizing cells from the same reference and predicted cells in the unannotated donor (Fig. 4A). Despite variations in tissue collection, staining protocols, imaging conditions, and segmentation workflows between these two donors—which can all affect marker intensities-CellFuse produced cell type annotations that closely matched the spatial structure and marker distribution in the new dataset (Fig. 4D, left). The predicted cell types were consistent with expected tissue architecture and known distributions of major intestinal cell types (Fig. 4D, right). To further validate CellFuse’s predictions, we examined the average marker expression profiles of predicted cell types in the unannotated donor and compared them to the expert-annotated data used as reference (Supplementary Fig. 6B). Hierarchical clustering of these profiles revealed close grouping of matched cell types across donors, further validating the reproducibility and biological consistency of CellFuse’s predictions (Supplementary Fig. 6B).

Finally, we assessed CellFuse’s performance in resolving complex tumor microenvironments (TMEs) using highly multiplexed IMC data from 12 breast cancer patients^19^. This dataset presented substantial inter-patient heterogeneity in tumor, stromal, and immune cell composition, offering a rigorous test of CellFuse’s ability to generalize across diverse clinical samples. We used annotated cells from four patients (n = 93,607) as a reference to predict cell types in the remaining eight patients (n = 286,647) (Fig. 4E). CellFuse successfully identified major cellular compartments within the TME including malignant epithelial cells, multiple immune subsets, and endothelial cells-achieving high prediction accuracy (F1 score = 0.93) (Fig. 4F and Supplementary Fig. 7A). Importantly, CellFuse preserved the spatial complexity of the TME across patients. It accurately captured distinct architectural patterns, such as “compartmentalized tumors” with spatially segregated immune and tumor regions (e.g., patient TBB129), and “immune-desert” phenotypes marked by sparse immune infiltration (e.g., patient TBB102) (Fig. 4G). Spatial overlays of predicted cell types across additional patients further demonstrated strong concordance with expert annotations (Supplementary Fig. 7B), reinforcing CellFuse’s ability to recover biologically meaningful structure in situ. Together, these results highlight the robustness, scalability, and translational relevance of CellFuse for high-resolution, cross-patient cell type annotation in spatial proteomics studies.

### Reconstructing spatially resolved tumor microenvironments via single-cell integration

One of the key challenges in single-cell research is accurately matching cell types across different modalities^24^. Cross-modal integration—such as between single-cell suspension and spatial proteomics presents a powerful opportunity to unify complementary cellular information. Accurately aligning cell types between these modalities enables researchers to not only identify diverse immune and stromal populations within tissues but also investigate their spatial relationships and organization. We reasoned that CellFuse’s robust cross-modal integration could be applied to transfer cell type annotations from single-cell suspension data (reference) onto spatial data (query), thereby enabling the inference of spatial localization for well-characterized immune and stromal cell types within the tissue microenvironment. To demonstrate this, we utilized a colon carcinoma dataset profiled using CyTOF and multiplexed ion beam imaging by time of flight (MIBI-TOF) to study the metabolic profiles of immune cells and their spatial relationship with tumor cells. To ensure stringency and generalizability, we used both healthy PBMCs and healthy colon CyTOF data as reference datasets and transferred their annotated cell type labels onto the spatial MIBI-TOF colon carcinoma dataset (query; Fig. 5A).

**Figure 5:**
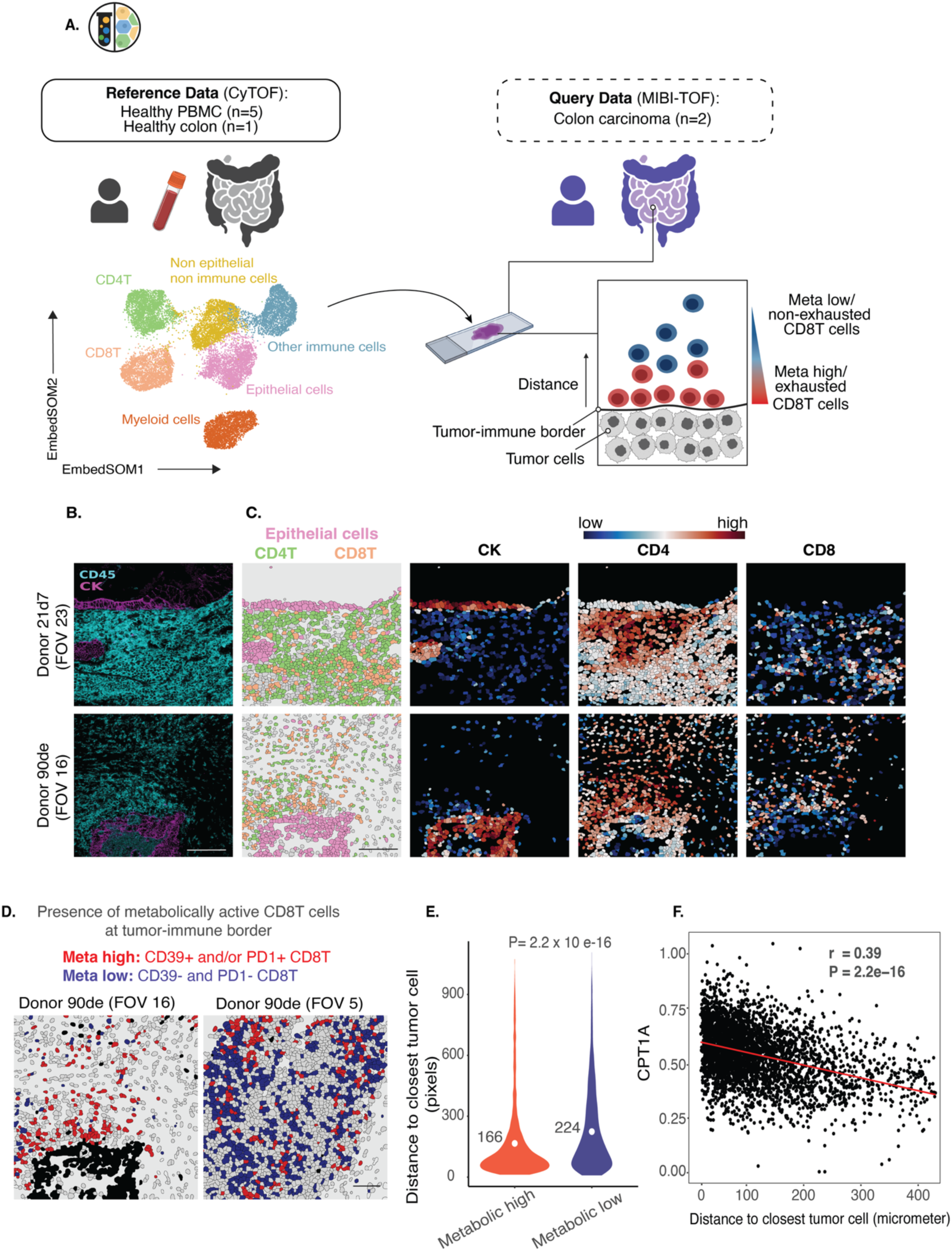
CellFuse maps tumor microenvironment by linking spatial and suspension data. **A.** Schematic overview of the Hartmann et al. CyTOF and MIBI-TOF datasets analyzed with CellFuse. The CellFuse model was trained on CyTOF data from healthy PBMCs and healthy colon, and used to predict cells in MIBI-TOF images from colon carcinoma patients and healthy colon tissue. **B.** Fluorescent images from donor 21d7 (top) and 90de (bottom). Markers are color-coded as follows: CK (cynan) and CD45 (blue). scale bar, 100 μm. **C.** Segmented images from donors 21d7 (top) and 90de (bottom), showing (left to right): CellFuse-predicted cell types, and expression levels of CK, CD4, and CD8 markers. scale bar, 100 μm. **D.** Two subset of CellFuse predicted CD8T cells (Meta high and Meta low) were visualized in the original images. scale bar, 100 μm. **E.** Two-sided Wilcoxon rank-sum test comparing the distance to the nearest tumor cell between Meta-high and Meta-low CD8⁺ T cells. Mean distances are indicated. **F.** Linear regression analysis reveals a significant inverse relationship between normalized asinh expression of metabolic enzyme (CPT1A) and distance to the closest tumor cell (P = 2.2 × 10⁻^16^). The red line represents the fitted regression model. CyTOF, Cytometry by Time of Flight; MIBI-TOF, Multiplexed Ion Beam Imaging by Time-Of-Flight; PBMC, Peripheral blood mononuclear cells; FOV; Field of View; CD8T, CD8 T cells; CD4T, CD4 T cells; CK, Cytokeratin; CPT1A, Carnitine Palmitoyltransferase 1A.

We first visualized the original multiplexed image (Fig. 5B) to define the tumor–immune boundary, using cytokeratin (CK, magenta) to mark tumor cells and CD45 (cyan) for immune cells. We then overlaid cell type annotations generated by CellFuse onto segmented single cells (Fig. 5C). The annotated immune (CD4T and CD8T cells) and tumor cell aligned well with the spatial organization observed in the raw image, demonstrating CellFuse’s ability to faithfully recapitulate spatially resolved tissue architecture. One key finding of the original study data was that CD8T cells positioned at the tumor–immune border exhibited markers of exhaustion, marked by CD39+ and/or PD-1+ expression, and showed elevated levels of metabolic markers compared to CD8T cells located further away from the boundary (Fig. 5A). We refer to these cells as Meta-high (border-proximal) and Meta-low (distal) CD8T cells, respectively. We investigated these CD8T cells using CellFuse-predicted annotations and observed a similar pattern that Meta-low CD39+/PD1+ CD8T cells were located farther from the tumor border compared to their Meta-high counterparts (Fig. 5D and E). Moreover, metabolic marker expression in these cells was strongly linked to their proximity to tumor cells, with cells closer to the tumor displaying higher metabolic activity—most notably marked by elevated carnitine palmitoyltransferase (CPT1A) expression (Fig. 5F). Together, these results demonstrate that CellFuse not only preserves spatial fidelity in its annotations but also enables biologically meaningful insights into the spatial-metabolic organization of immune cells within the tumor microenvironment.

## Methods

### The CellFuse workflow

CellFuse accepts as input two datasets: a labeled reference dataset (e.g., CyTOF or CITE-seq with known cell type labels) and an unlabeled query dataset (e.g., CITE-seq, CODEX, or IMC data without labels). Both datasets are represented as matrices of protein or gene expression profiles, where rows represent cells and columns represent shared features (e.g., proteins). Let 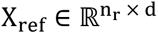 be the reference dataset with n_r_ cells and d shared features and let 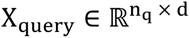 be the query dataset. The reference dataset is first split into training and validation fractions and is accompanied by a matrix of one-hot encoded cell type labels 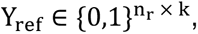 where k is the number of cell types.

#### Stage 1: Model Training

CellFuse uses a multi-layer perceptron (MLP) encoder f_θ_ to project input features into a lower-dimensional embedding space. Formally, for an input cell X ∈ ℝ^d^ the encoder outputs an embedding:

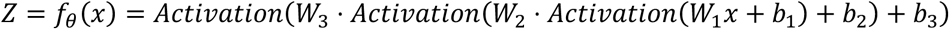

where *W₁, W₂, W₃* and *b₁, b₂, b₃* are learnable parameters. The non-linear activation function is user-configurable and can be set to ReLU, ELU, or LeakyReLU. Batch normalization and dropout are optionally applied for regularization.

Training steps:

1. Data Input: Two labeled datasets are provided for training and validation, enabling performance monitoring and early stopping.
2. Data Normalization: Each dataset is independently z-score normalized across all features to ensure consistent scaling.
3. Label Encoding: The cell type annotations in the training set are one-hot encoded into a matrix 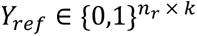 where k is the number of cell types. These labels are used for supervised training and contrastive loss calculation.
4. Representation Learning: The MLP encoder maps cells into a shared embedding space where cells of the same type cluster together. The activation function (ReLU, ELU, or LeakyReLU) is chosen by the user based on data properties.
5. Contrastive Loss: A supervised contrastive loss ℒ_contrastive_ is minimized to bring embeddings of the same label closer while pushing apart those from different classes:

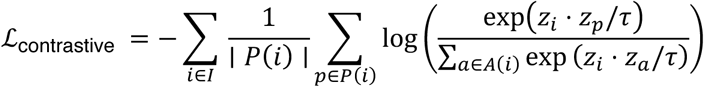

where:

- *z_i_* is the embedding of anchor cell iii,
- *P*(*i*) denotes the set of positives (same class),
- *A*(*i*) includes all anchors except iii,
- *τ* is a temperature scaling parameter.
6. **Optimization and Early Stopping**: Training is performed using the Adam optimizer with optional weight decay. Model performance is monitored on the user-provided validation set after each epoch. Early stopping is triggered if the validation loss does not improve within a specified patience window, preventing overfitting.

#### Stage 2: Predicting Cells

After training, the encoder *f*_*θ*_ is applied to both reference and query datasets to obtain embeddings *Z_ref_* and *Z_query_*. A k-nearest neighbor (kNN) classifier is trained on ℤ*_ref_* using the known labels *Y_ref_* and used to predict labels for *Z_query_*. Each predicted cell is also assigned a confidence score based on the softmax-normalized similarity to its neighbors. Formally, given the embedding 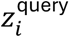 of a query cell, we identify its k nearest neighbors among the reference embeddings and assign a label based on majority vote or weighted confidence:

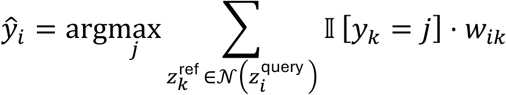

Where *w_ik_* denotes the similarity weight between 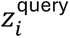 and 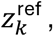 and 𝕀 is the indicator function.

#### Stage 3: Data Integration

For each predicted cell type c, the reference data 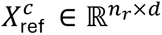 is used to perform principal component analysis (PCA):

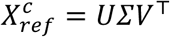

This yields a rotation matrix *V* ∈ ℝ*d* × *d*, which defines a reference-specific embedding space. The corresponding query data 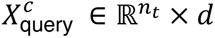 is projected into this PCA space using:

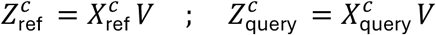

Step 2: Distribution Alignment in PCA Space

To harmonize the distributions of query embeddings with those of the reference, each principal component of the query data is aligned to the corresponding reference component using empirical distribution matching. This is achieved by reordering the query values based on the ranked structure of the reference:

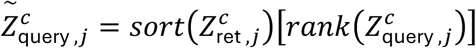

If the number of reference and query cells differ, the reference component values are resampled (with or without replacement) to match the size of the query distribution. Zero-valued entries are retained from the original input to preserve sparsity.

Step 3: Reconstruction into Original Feature Space

The aligned query embeddings 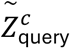 are then projected back to the original feature space using the transpose of the PCA rotation matrix:

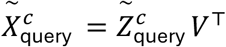

This yields a corrected query matrix 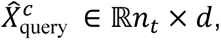 in which feature distributions are aligned to the reference while maintaining the biological structure inferred from the query data.

### Benchmarks

#### Integration methods

We evaluated the integration performance of CellFuse in comparison to three widely used tools: Seurat (v5.2.1)^10^ ^11^, Harmony (v1.2.3), and FastMNN (from the *batchelor* R package, v1.22.0)^12^. All methods were executed using their default parameters.

#### Spatial cell type prediction methods

To assess cell type prediction accuracy in spatial proteomics data, we compared CellFuse to three baseline methods: CELESTA (v 0.0.0.9), Astir (v 0.1.5), SVM implemented via the e1071 package (v1.7-16)^22^, and Seurat (v5.2.1)^10^. All models were trained and evaluated using default settings.

##### Metrics

To evaluate the quality of cross-modality data integration, we employed a suite of metrics from the scIB benchmarking framework as described by Luecken et al. ^16^. These metrics assess two key integration objectives: the preservation of biological identity and the effective mixing of data across batches or modalities.

Biological conservation was assessed using the following metrics:

Cell type Average Silhouette Width (ASW): quantifies the cohesion of cells within the same cell type and their separation from other types. Higher values indicate better preservation of biological structure.

Normalized Mutual Information (NMI) and Adjusted Rand Index (ARI): compare clustering results from the integrated data to ground truth cell type annotations, with higher scores indicating better retention of biological identity post-integration.

Batch effect correction was evaluated using:

Graph iLISI (inverse Local Inverse Simpson’s Index): measures the diversity of batch labels in each cell’s local neighborhood within a k-nearest neighbor graph. A higher iLISI score reflects effective local batch mixing and reduced modality-specific bias.

Batch ASW: assesses how well cells from different batches are intermixed within a given cell type. Higher batch ASW values suggest successful integration across batches or modalities.

Graph Connectivity: quantifies the continuity of the shared nearest neighbor graph across batches. High connectivity indicates successful alignment of cell populations across datasets.

To summarize overall performance, we computed a composite integration score as a weighted average of the mean biological conservation score (weight = 0.6) and the mean batch mixing score (weight = 0.4), following established recommendations^16^. Full definitions and implementation details of these metrics are provided in the scIB package documentation and reference therein.

##### Datasets

###### Bone Marrow (BM) Dataset

This dataset contains healthy BM cells profiled using both CyTOF and CITE-seq modalities. The CyTOF dataset includes 265,627 cells measured with a 32-marker antibody panel as described by Levine et al.^13^, while the CITE-seq dataset comprises 30,672 cells profiled with a 25-marker antibody panel reported by Stuart et al^10^. Both modalities share 12 overlapping protein markers and 8 common cell types. For benchmarking purposes, 20,000 cells were randomly subsampled using the tof_downsample_constant() function from the TidyTOF R package.

###### Peripheral Blood Mononuclear Cells (PBMC) Dataset

This dataset consists of PBMCs collected from healthy donors and measured using a 45-marker CyTOF panel^14^ and a 228-marker CITE-seq panel^15^. The two datasets share 27 overlapping protein markers and cover 12 common immune cell types, supporting robust evaluation of cross-modality integration. Both datasets were downsampled to 40,000 cells using the tof_downsample_constant() function from the TidyTOF package for benchmarking^26^.

###### CAR-T Cell Therapy Dataset

This dataset includes published single-cell CyTOF and CITE-seq data from 28 patients with large B-cell lymphoma (LBCL) who underwent axi-cel CAR-T cell therapy^17^. Peripheral blood mononuclear cells were collected at pre- and post-infusion timepoints. The dataset was retrieved from Good et al. (accession number: GSE168940).

###### HuBMAP CODEX Intestine Dataset

This dataset comprises CODEX images of healthy human intestine acquired using a 48-marker antibody panel, as reported by Brbic et al^27^. Expert-annotated data from one donor includes 248,285 cells spanning 21 distinct cell types, while an unannotated donor dataset includes 619,186 cells.

###### Breast Cancer IMC Dataset

This dataset includes Imaging Mass Cytometry (IMC) data from breast tumor tissue samples of 12 patients. The dataset was generated as part of the study by Tietscher et al^19^. and provides cell type annotations for 10 distinct immune and stromal populations.

###### Colon Cancer CyTOF and MIBI-TOF Dataset

This dataset was generated by Hartmann et al.^19, 28^ and consists of 71,789 cells profiled by CyTOF from healthy peripheral blood and colon carcinoma patients, annotated into six major cell types. Additionally, MIBI-TOF imaging data from two colon cancer patients, comprising 63,747 cells, were used to evaluate CellFuse’s performance in spatial integration and annotation.

## Discussion

CellFuse is a deep learning-based framework for cross-modality single-cell data integration, specifically designed to remain robust even in cases where feature overlap between datasets is minimal. It addresses a central challenge in the field: integrating single-cell proteomics data across platforms that often measure non-overlapping or weakly overlapping protein panels. This challenge is particularly acute when aligning spatial proteomic datasets (e.g., CODEX, IMC) with suspension-based references such as CyTOF or CITE-seq, where the number of shared markers is limited and often noisy^29^. In these scenarios, traditional anchor- or graph-based methods often fail due to the sparse or noisy nature of shared features.

The CellFuse pipeline incorporates several innovations that make it particularly robust to such weakly linked conditions. It employs supervised contrastive learning to train an encoder on labeled reference data, learning a shared latent space in which biologically similar cells are embedded close together, regardless of their originating modality. Query cells are then projected into this space for both integration and cell type prediction. By capturing both shared and modality-specific information, CellFuse remains resilient to challenges such as marker dropout, imbalanced feature panels, and partial cell type coverage conditions that frequently arise in translational or clinical datasets.

We demonstrate that CellFuse enables highly accurate cross-modal cell type annotation, consistently outperforming existing methods in settings where feature linkage is weak. For example, CellFuse accurately transferred annotations from CyTOF references to spatial proteomic datasets, resolving fine-grained cell types in complex tissues such as human intestine and breast tumors. It also enabled the identification of clinically important CAR-T cell subpopulations in feature-limited CITE-seq query data. These results highlight CellFuse’s ability to maintain biological resolution even under challenging integration conditions.

To evaluate CellFuse in the context of existing tools, we benchmarked its performance against Seurat^10^, Harmony^11^, and FastMNN^12^ -three widely used methods for single-cell integration. While these methods are effective in well-matched transcriptomic datasets with high feature overlap, they show significant performance degradation in cross-modality settings. Specifically, Seurat relies on anchor-based transfer, which fails when shared features are limited or uninformative. Harmony is effective for batch correction but is not modality-aware, often collapsing rare populations or blurring distinctions between closely related cell types. FastMNN, which uses mutual nearest neighbor matching, is highly sensitive to population imbalance and performs poorly in heterogeneous datasets with modality mismatch.

In contrast, CellFuse consistently achieved higher annotation accuracy and better preservation of biological structure across a range of weak-linkage settings. For example, in a sequential feature dropping benchmark, CellFuse maintained high F1 scores even after most overlapping markers were removed, while competing methods failed to identify cell types beyond a critical threshold. These results demonstrate that CellFuse is more robust to decreasing feature overlap, a common limitation in practical multi-modal studies.

However, CellFuse is not without limitations. Its performance depends on the quality and granularity of the labeled reference dataset; errors or bias in the reference can propagate through the learned embedding space. Furthermore, its supervised nature may limit its ability to discover novel cell types that are not represented in the training data. In contrast, unsupervised or semi-supervised models like scANVI^30^ and SpaGCN^31^ can identify new clusters or incorporate spatial context more directly—capabilities not currently built into CellFuse. Additionally, CellFuse does not explicitly model cell-cell interactions or developmental trajectories, which may be important in some applications such as developmental biology or tumor microenvironment analysis.

Despite these caveats, CellFuse offers a scalable, efficient, and generalizable solution for integrating single-cell datasets across technologies. It supports rapid inference, is compatible with pre-trained models, and can be extended to new data types and applications. These features position CellFuse as a valuable addition to the integration toolkit, particularly for researchers working with heterogeneous, noisy, or clinically derived datasets where conventional methods fall short.

## Code and Data availability

The software package implementing the CellFuse algorithm, datasets analyzed in this manuscript and corresponding analysis scripts will be made public upon acceptance.

## Author contributions

A.K. and K.L.D conceptualized the study. A.K. was responsible for algorithm development, implementation, code development, and data analysis. A.K, S.C.B., Z.D, K.L.D. the data. A.K. generated the figures. S.R.V. tested the implemented algorithm. A.K, Z.D, S.C.B., K.L.D wrote and edited the manuscript. K.L.D. provided guidance, scientific input, and obtained funding.

## Acknowledgments

We would like to thank all members of the Davis laboratory and Pablo Domizi for helpful discussions. This work was supported by the National Institutes of Health R01-CA251858, the Mark Foundation ASPIRE award, and the Stanford Maternal Child Health Research Institute, MCHRI. A.K. is supported by the International Society for the Advancement of Cytometry (ISAC)’s Marylou Ingram Scholars program. K.L.D is supported by the Anne T. and Robert M. Bass Endowed Faculty Scholar in Pediatric Cancer and Blood Disease and the Harriet and Mary Zelencik Endowment.

## References

1. Baysoy, A., Bai, Z., Satija, R. & Fan, R. The technological landscape and applications of single-cell multi-omics. Nat. Rev. Mol. Cell Biol. 24, 695–713 (2023).

2. Baumgarth, N. & Roederer, M. A practical approach to multicolor flow cytometry for immunophenotyping. J. Immunol. Methods 243, 77–97 (2000).

3. Koladiya, A. & Davis, K. L. Advances in Clinical Mass Cytometry. Clin. Lab. Med. 43, 507– 519 (2023).

4. Konecny, A. J., Mage, P. L., Tyznik, A. J., Prlic, M. & Mair, F. OMIP-102: 50-color phenotyping of the human immune system with in-depth assessment of T cells and dendritic cells. Cytometry A. 105, 430–436 (2024).

5. Mair, F. et al. A Targeted Multi-omic Analysis Approach Measures Protein Expression and Low-Abundance Transcripts on the Single-Cell Level. Cell. Rep. 31, 107499 (2020).

6. Zhang, X. et al. An immunophenotype-coupled transcriptomic atlas of human hematopoietic progenitors. Nat. Immunol. 25, 703–715 (2024).

7. Goltsev, Y. et al. Deep Profiling of Mouse Splenic Architecture with CODEX Multiplexed Imaging. Cell 174, 968–981.e15 (2018).

8. Angelo, M. et al. Multiplexed ion beam imaging of human breast tumors. Nat. Med. 20, 436– 442 (2014).

9. Giesen, C. et al. Highly multiplexed imaging of tumor tissues with subcellular resolution by mass cytometry. Nat. Methods 11, 417–422 (2014).

10. Stuart, T. et al. Comprehensive Integration of Single-Cell Data. Cell 177, 1888–1902.e21 (2019).

11. Korsunsky, I. et al. Fast, sensitive and accurate integration of single-cell data with Harmony. Nat. Methods 16, 1289–1296 (2019).

12. Haghverdi, L., Lun, A. T. L., Morgan, M. D. & Marioni, J. C. Batch effects in single-cell RNA-sequencing data are corrected by matching mutual nearest neighbors. Nat. Biotechnol. 36, 421– 427 (2018).

13. Levine, J. H. et al. Data-Driven Phenotypic Dissection of AML Reveals Progenitor-like Cells that Correlate with Prognosis. Cell 162, 184–197 (2015).

14. COvid-19 Multi-omics Blood ATlas (COMBAT) Consortium. Electronic address: & COvid-19 Multi-omics Blood ATlas (COMBAT) Consortium. A blood atlas of COVID-19 defines hallmarks of disease severity and specificity. Cell 185, 916–938.e58 (2022).

15. Hao, Y. et al. Integrated analysis of multimodal single-cell data. Cell 184, 3573–3587.e29 (2021).

16. Luecken, M. D. et al. Benchmarking atlas-level data integration in single-cell genomics. Nat. Methods 19, 41–50 (2022).

17. Good, Z. et al. Post-infusion CAR T(Reg) cells identify patients resistant to CD19-CAR therapy. Nat. Med. 28, 1860–1871 (2022).

18. Hickey, J. W. et al. Organization of the human intestine at single-cell resolution. Nature 619, 572–584 (2023).

19. Tietscher, S. et al. A comprehensive single-cell map of T cell exhaustion-associated immune environments in human breast cancer. Nat. Commun. 14, 98–w (2023).

20. Zhang, W. et al. Identification of cell types in multiplexed in situ images by combining protein expression and spatial information using CELESTA. Nat. Methods 19, 759–769 (2022).

21. Geuenich, M. J. et al. Automated assignment of cell identity from single-cell multiplexed imaging and proteomic data. Cell. Syst. 12, 1173–1186.e5 (2021).

22. Cortes, C. & Vapnik, V. Support-vector networks. Mach. Learning 20, 273–297 (1995).

23. Huynh, K. L. A. et al. Deconvolution of cell types and states in spatial multiomics utilizing TACIT. Nat. Commun. 16, 3747–4 (2025).

24. Lance, C., et al. Multimodal single cell data integration challenge: results and lessons learned. bioRxiv, 2022.04.11.487796 (2022).

25. Keyes, T. J., Koladiya, A., Lo, Y., Nolan, G. P. & Davis, K. L. tidytof: a user-friendly framework for scalable and reproducible high-dimensional cytometry data analysis. Bioinformatics Advances 3, vbad071 (2023).

26. Brbić, M. et al. Annotation of spatially resolved single-cell data with STELLAR. Nat. Methods 19, 1411–1418 (2022).

27. Hartmann, F. J. et al. Single-cell metabolic profiling of human cytotoxic T cells. Nat. Biotechnol. 39, 186–197 (2021).

28. Confident multimodal analysis of single cells across platforms and speciesNature Methods 20, 191–192 (2023).

29. Xu, C. et al. Probabilistic harmonization and annotation of single-cell transcriptomics data with deep generative models. Mol. Syst. Biol. 17, e9620 (2021).

30. Hu, J. et al. SpaGCN: Integrating gene expression, spatial location and histology to identify spatial domains and spatially variable genes by graph convolutional network. Nat. Methods 18, 1342–1351 (2021).

