## Supplementary Figures 1-7 for "CellFuse Enables Multi-modal Integration of Single-cell and Spatial Proteomics data"

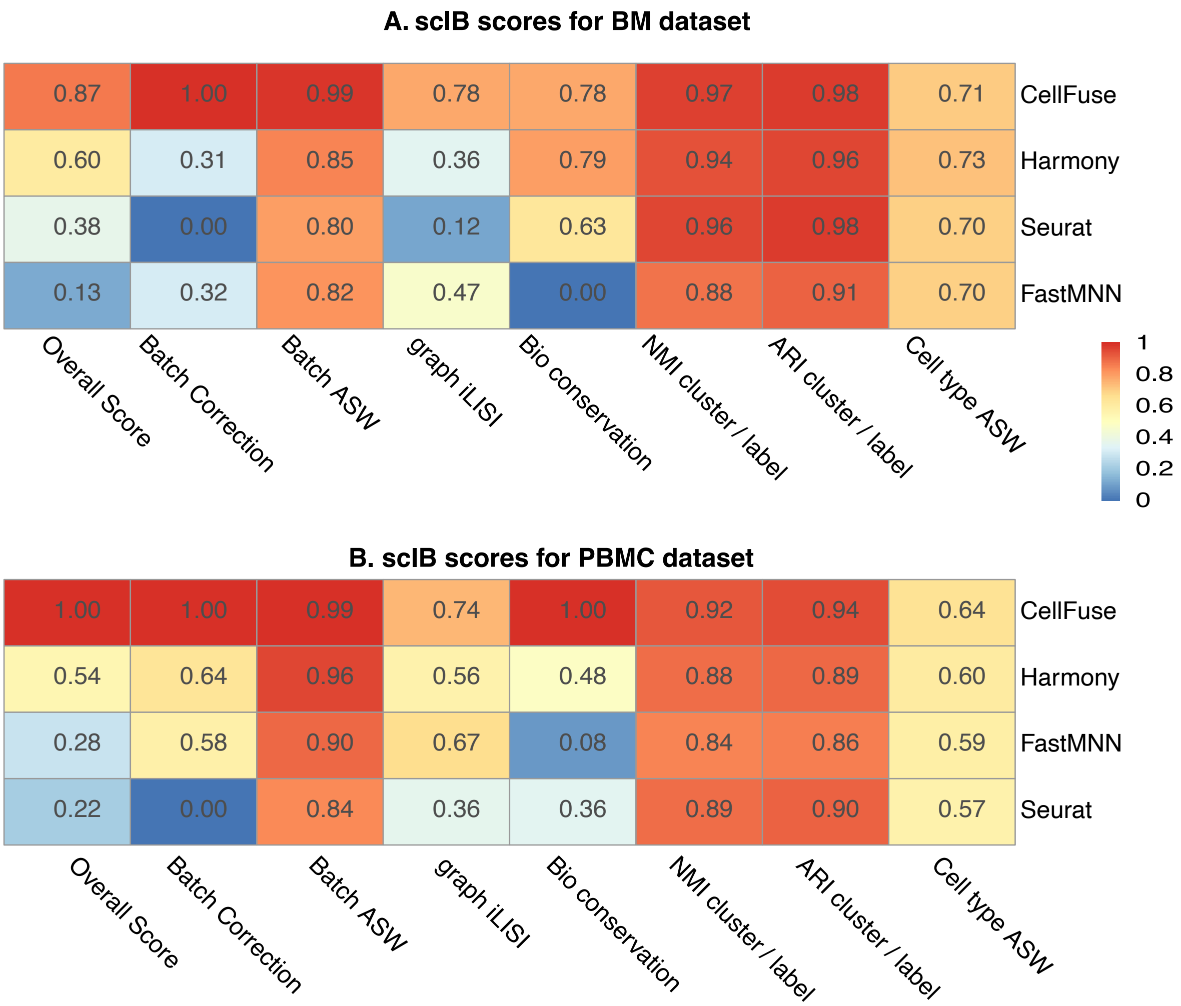

**Supplemental Figure 1: scIB benchmarking scores for (A) BM and (B) PBMC dataset across integration methods.**

The heatmap summarizes seven integration metrics for CellFuse, Harmony, FastMNN, and Seurat. Metrics include batch correction (Batch ASW, graph iLISI), biological conservation (NMI cluster/label, ARI cluster/label, Cell type ASW), and an overall integration score.

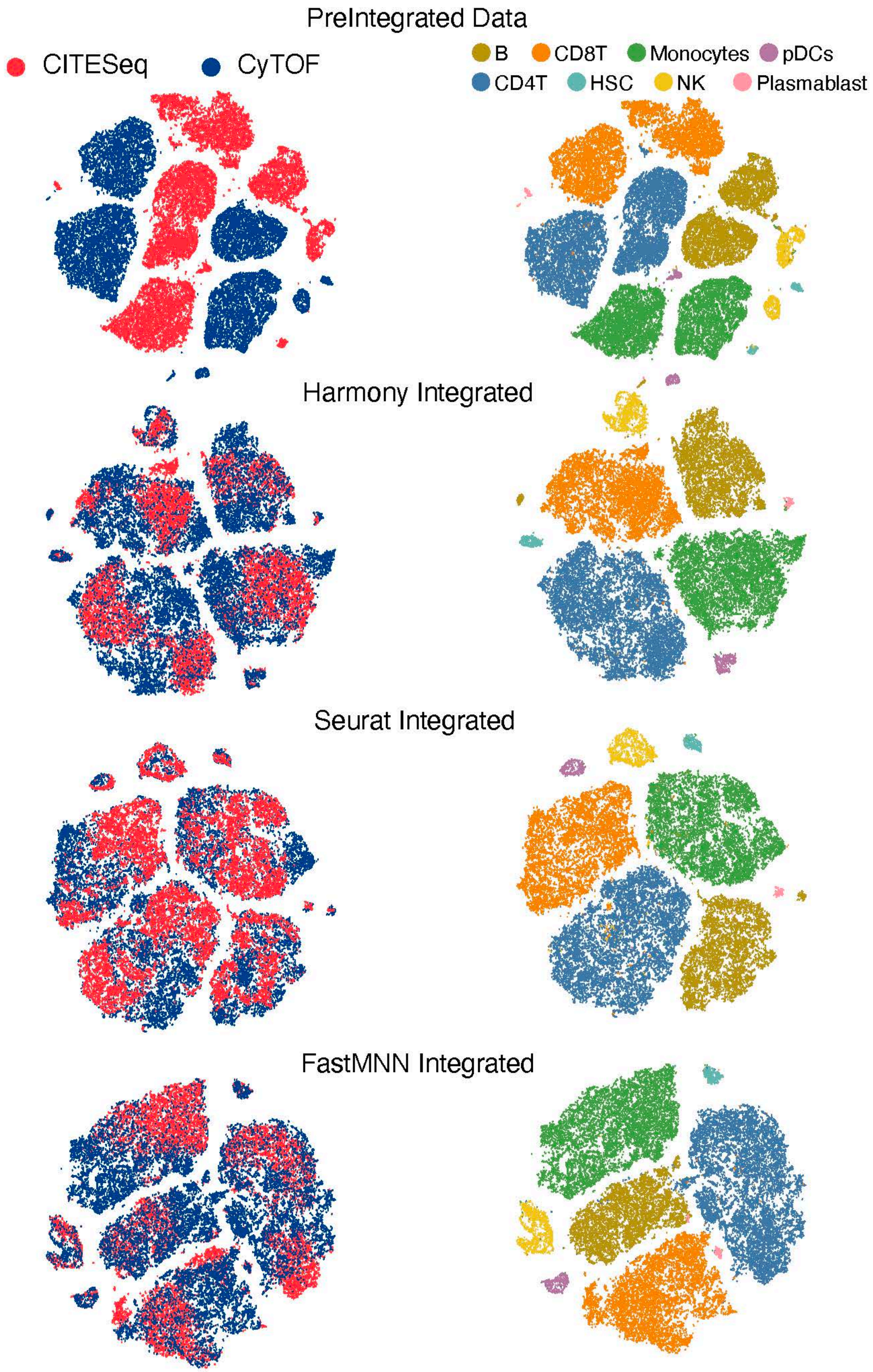
PreIntegrated Data
CITESeq
CyTOF
B
CD8T
Monocytes
pDCs
CD4T
HSC
NK
Plasmablast
Harmony Integrated
Seurat Integrated
FastMNN Integrated

**Supplemental Figure 2: tSNE visualization of pre and post-integration of BM CITESeq and CyTOF data**

t-SNE plots visualizing pre-integration and post-integration results with the various methods. All cells are colored by modalities (left) and cell type annotations (right).

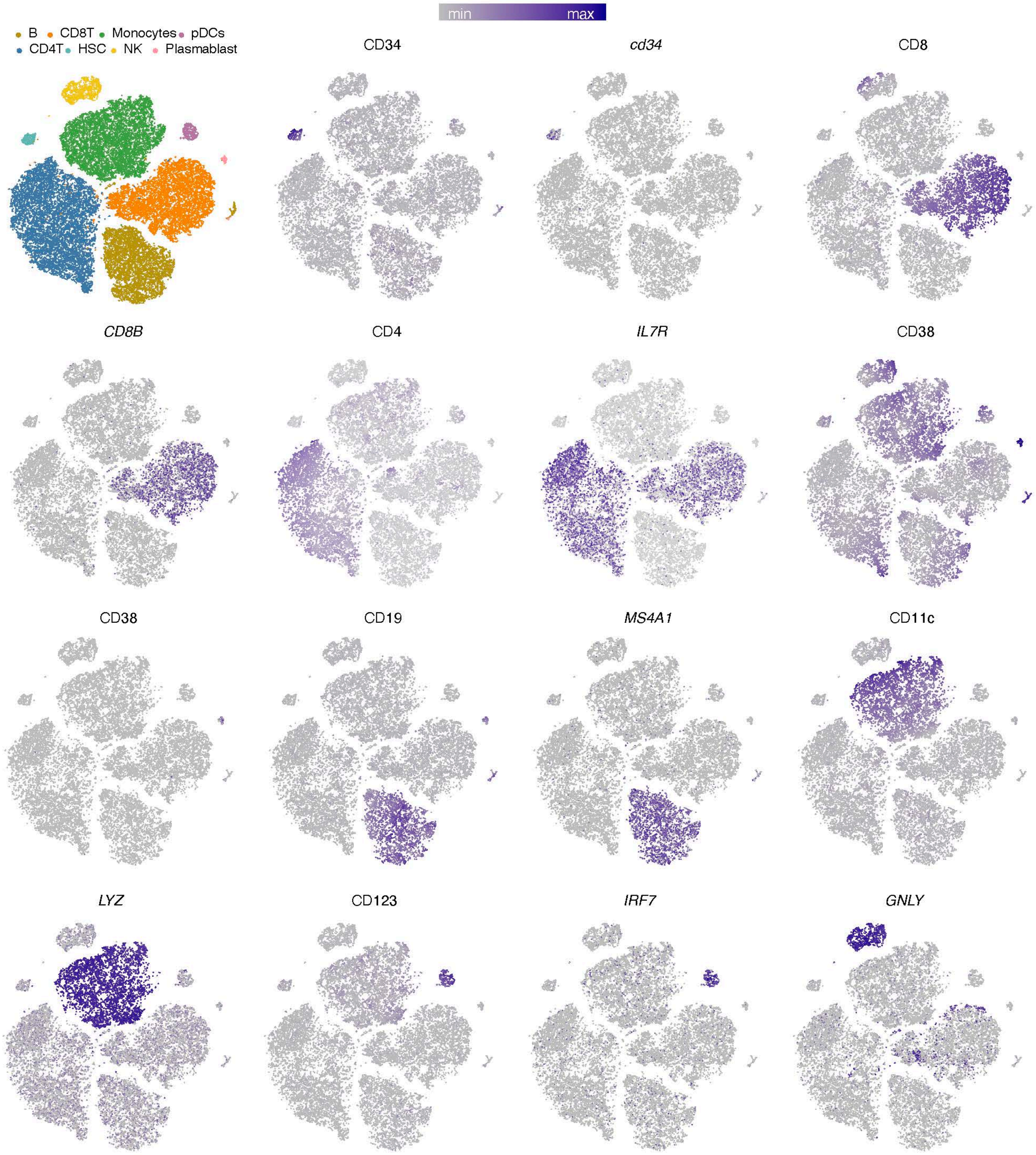
min
max
B
CD8T
Monocytes
pDCs
CD4T
HSC
NK
Plasmablast
CD34
cd34
CD8
CD8B
CD4
IL7R
CD38
CD38
CD19
MS4A1
CD11c
LYZ
CD123
IRF7
GNLY

**Supplemental Fig 3: tSNE visualization of CellFuse integrated BM CITESeq and CyTOF data**

**A.** CellFuse integrated tSNE plot colored by cell types

**B.** t-SNE plot showing *protein* and *transcript* expression profiles of CellFuse integrated BM data. Plots depicting *transcript* expression are titled in italics.

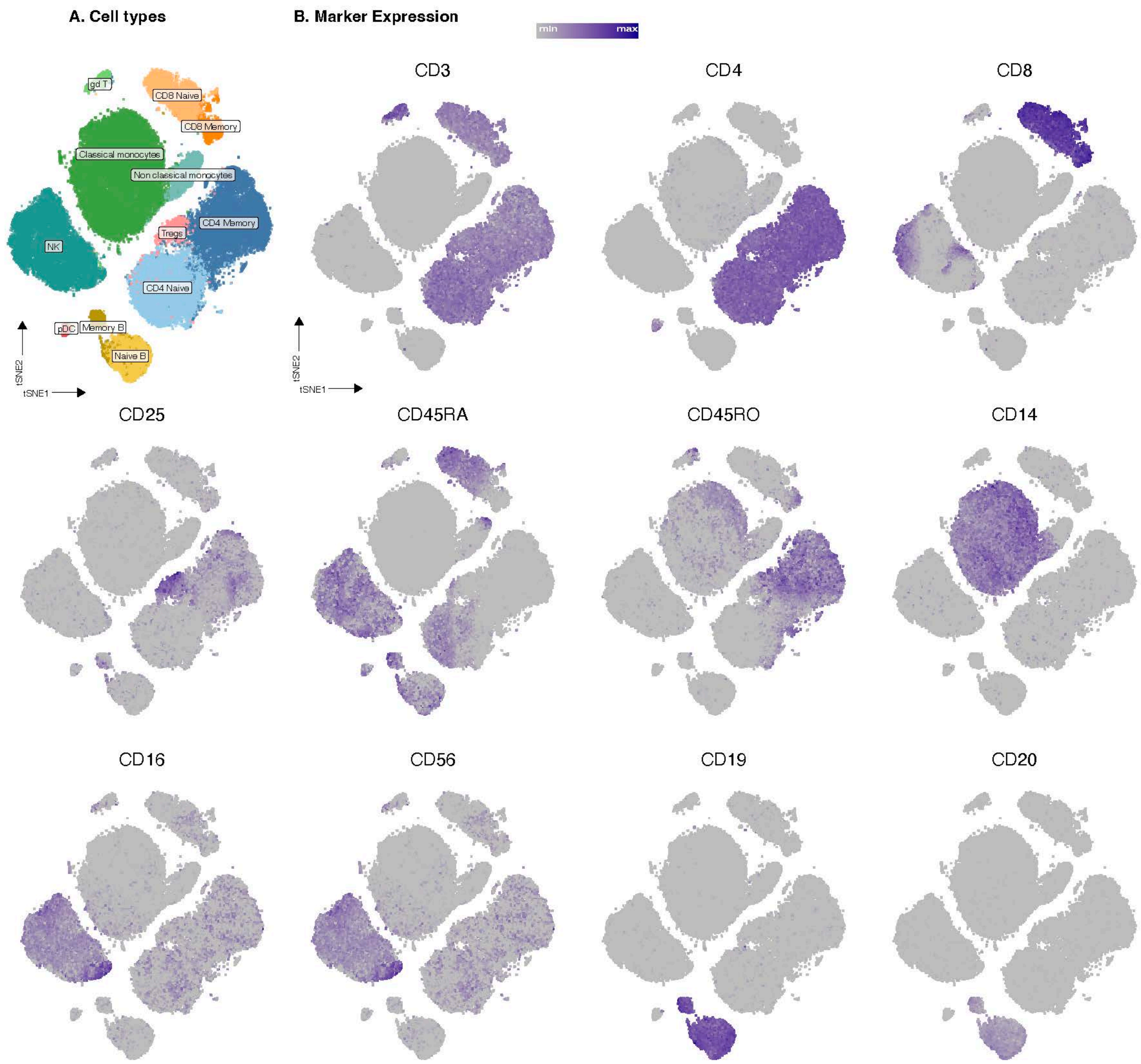

A. Cell types
B. Marker Expression
min
max
gd T
CD8 Naive
CD8 Memory
Classical monocytes
Non classical monocytes
CD4 Memory
Tregs
NK
CD4 Naive
pDC
Memory B
Naive B
tSNE2
tSNE1
CD3
CD4
CD8
CD25
CD45RA
CD45RO
CD14
CD16
CD56
CD19
CD20

**Supplemental Fig 4: tSNE visualization of CellFuse integrated PBMC data**
**A.** CellFuse integrated tSNE plot colored by cell types

**B.** t-SNE plot showing *protein* expression of CellFuse integrated PBMC data.

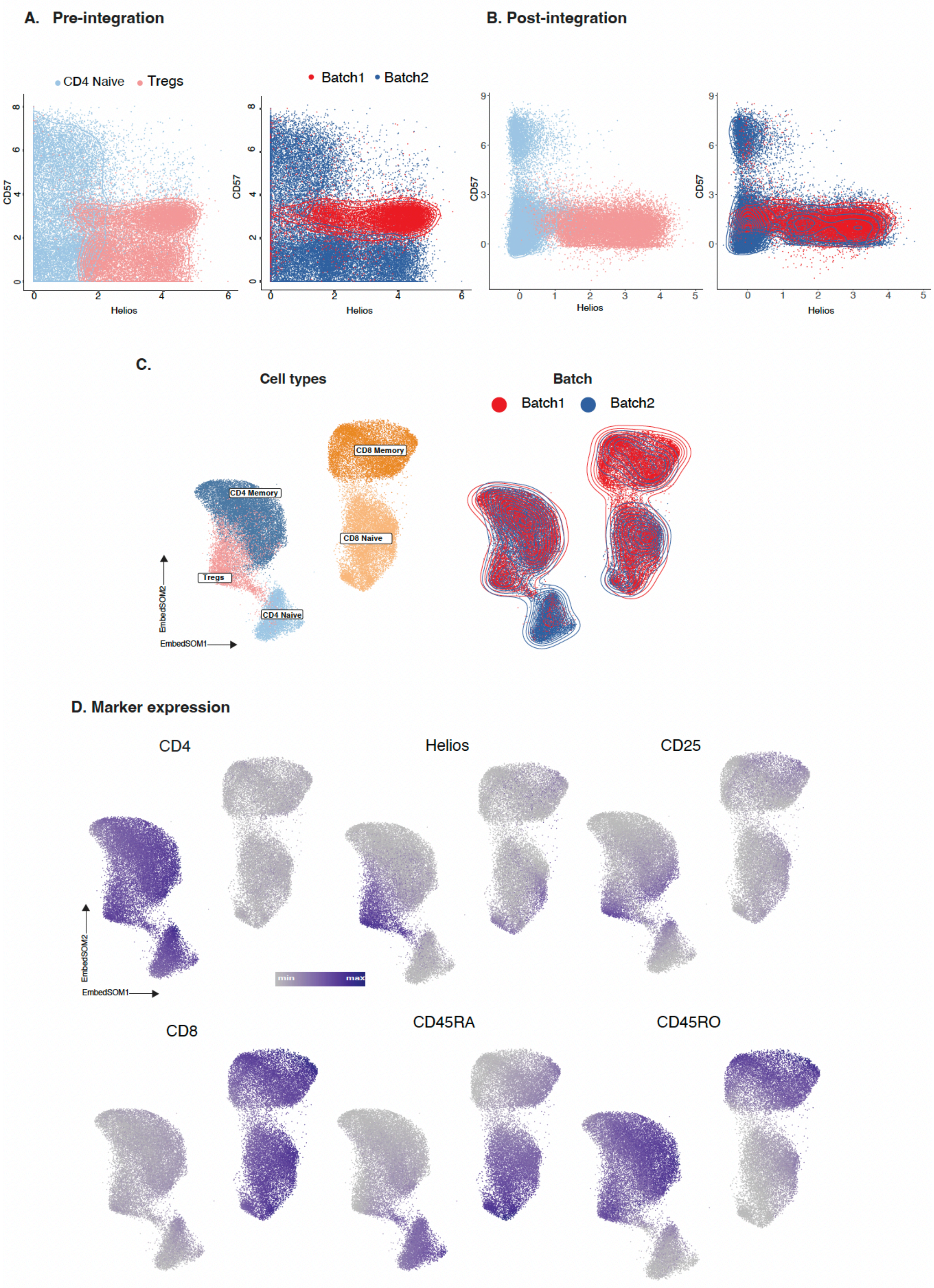
A. Pre-integration
B. Post-integration
CD4 Naive
Tregs
Batch1
Batch2
CD57
Helios
C.
Cell types
Batch
CD8 Memory
CD4 Memory
CD8 Naive
Tregs
CD4 Naive
EmbedSOM2
EmbedSOM1
D. Marker expression
CD4
Helios
CD25
min
max
CD8
CD45RA
CD45RO

**Supplemental Fig 5: CAR T cell analysis using CellFuse**

**A.** Batch effect detection in pre-integrated CD4 Naive and Tregs. Cells are colored by cell types (left) and by batches (right).

**B.** CellFuse integrated cells colored by cell types (left) and by batches (right).

**C.** EmbedSOM visualization of CellFuse integrated cells colored by cell types (left) and batches (right).

**D.** EmbedSOM plots colored by protein expression.

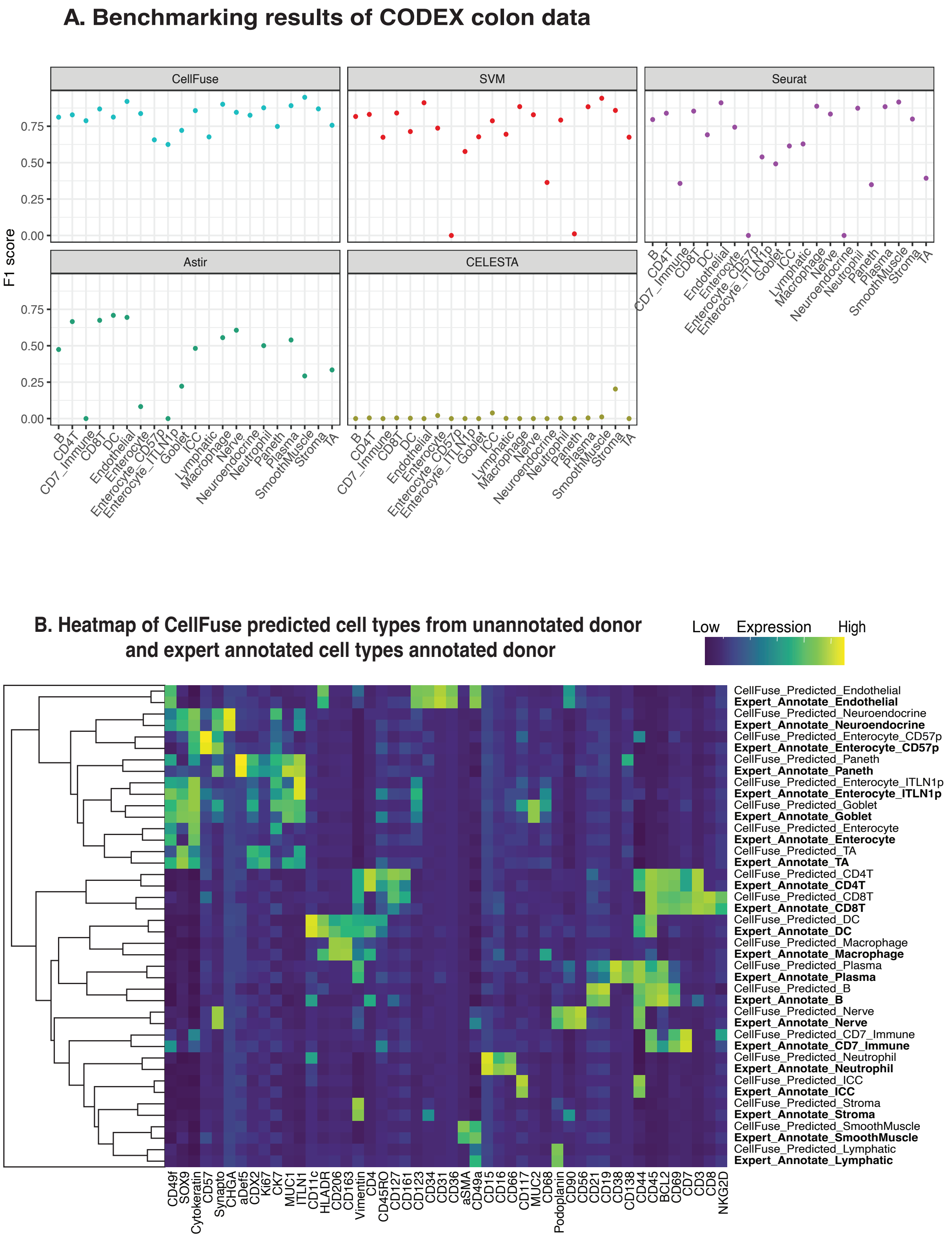
A. Benchmarking results of CODEX colon data
CellFuse
SVM
Seurat
Astir
CELESTA
F1 score
0.00
0.25
0.50
0.75
B
CD4T
CD7_Immune
CD8T
DC
Endothelial
Enterocyte
Enterocyte_CD57p
Enterocyte_ITLN1p
Goblet
ICC
Lymphatic
Macrophage
Nerve
Neuroendocrine
Neutrophil
Paneth
Plasma
SmoothMuscle
Stroma
TA
B. Heatmap of CellFuse predicted cell types from unannotated donor and expert annotated cell types annotated donor
Low Expression High
CellFuse_Predicted_Endothelial
Expert_Annotate_Endothelial
CellFuse_Predicted_Neuroendocrine
Expert_Annotate_Neuroendocrine
CellFuse_Predicted_Enterocyte_CD57p
Expert_Annotate_Enterocyte_CD57p
CellFuse_Predicted_Paneth
Expert_Annotate_Paneth
CellFuse_Predicted_Enterocyte_ITLN1p
Expert_Annotate_Enterocyte_ITLN1p
CellFuse_Predicted_Goblet
Expert_Annotate_Goblet
CellFuse_Predicted_Enterocyte
Expert_Annotate_Enterocyte
CellFuse_Predicted_TA
Expert_Annotate_TA
CellFuse_Predicted_CD4T
Expert_Annotate_CD4T
CellFuse_Predicted_CD8T
Expert_Annotate_CD8T
CellFuse_Predicted_DC
Expert_Annotate_DC
CellFuse_Predicted_Macrophage
Expert_Annotate_Macrophage
CellFuse_Predicted_Plasma
Expert_Annotate_Plasma
CellFuse_Predicted_B
Expert_Annotate_B
CellFuse_Predicted_Nerve
Expert_Annotate_Nerve
CellFuse_Predicted_CD7_Immune
Expert_Annotate_CD7_Immune
CellFuse_Predicted_Neutrophil
Expert_Annotate_Neutrophil
CellFuse_Predicted_ICC
Expert_Annotate_ICC
CellFuse_Predicted_Stroma
Expert_Annotate_Stroma
CellFuse_Predicted_SmoothMuscle
Expert_Annotate_SmoothMuscle
CellFuse_Predicted_Lymphatic
Expert_Annotate_Lymphatic
CD49f
SOX9
Cytokeratin
CD57
Synapto
CHGA
aDef5
CDX2
Ki67
CK7
MUC1
ITLN1
CD11c
HLADR
CD206
CD163
Vimentin
CD4
CD45RO
CD127
CD161
CD123
CD34
CD31
CD36
aSMA
CD49a
CD15
CD16
CD66
CD117
MUC2
CD68
Podoplanin
CD90
CD56
CD21
CD19
CD38
CD138
CD44
CD45
BCL2
CD69
CD7
CD3
CD8
NKG2D

**Supplemental Fig 6: Benchmarking and evaluation of CODEX HuBMAP data**

**A.** Performance evaluation of predicted cell types by CellFuse, SVM, Seurat, Astir and CELESTA using F1 score

**B.** Comparison of average marker expression for CellFuse predicted cell types in unannotated and expert-annotated donors. CellFuse predicted cell types are labeled in bold and italics.

SVM, Support Vector Machine; DC, dendritic cell; ICC, interstitial cells of Cajal; TA, transit amplifying cell.

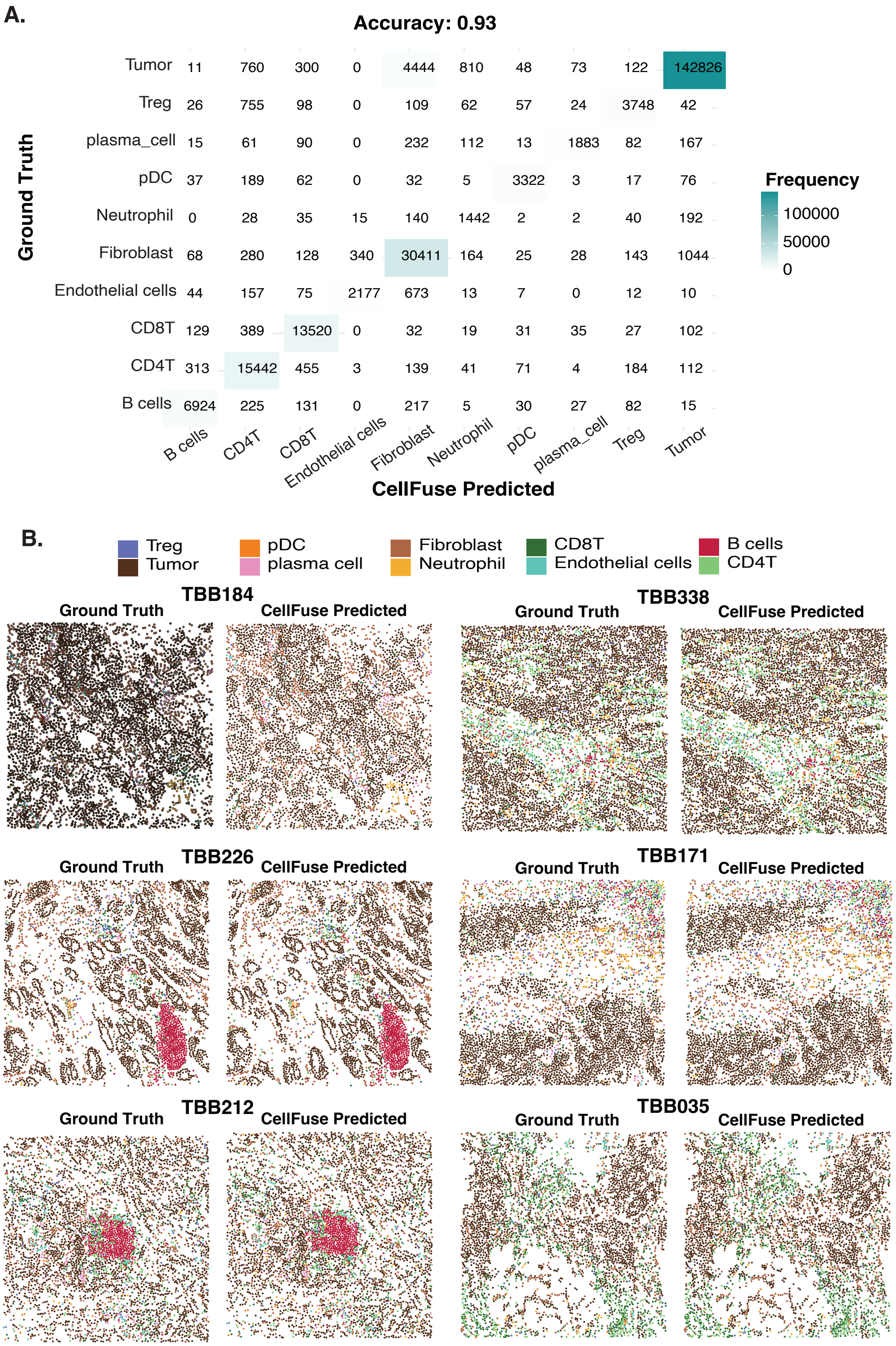
A.
Accuracy: 0.93
Ground Truth
Tumor 11 760 300 0 4444 810 48 73 122 142826
Treg 26 755 98 0 109 62 57 24 3748 42
plasma_cell 15 61 90 0 232 112 13 1883 82 167
pDC 37 189 62 0 32 5 3322 3 17 76
Neutrophil 0 28 35 15 140 1442 2 2 40 192
Fibroblast 68 280 128 340 30411 164 25 28 143 1044
Endothelial cells 44 157 75 2177 673 13 7 0 12 10
CD8T 129 389 13520 0 32 19 31 35 27 102
CD4T 313 15442 455 3 139 41 71 4 184 112
B cells 6924 225 131 0 217 5 30 27 82 15
B cells CD4T CD8T Endothelial cells Fibroblast Neutrophil pDC plasma_cell Treg Tumor
CellFuse Predicted
Frequency
100000
50000
0
B.
Treg
Tumor
pDC
plasma cell
Fibroblast
Neutrophil
CD8T
Endothelial cells
B cells
CD4T
TBB184
Ground Truth
CellFuse Predicted
TBB338
Ground Truth
CellFuse Predicted
TBB226
Ground Truth
CellFuse Predicted
TBB171
Ground Truth
CellFuse Predicted
TBB212
Ground Truth
CellFuse Predicted
TBB035
Ground Truth
CellFuse Predicted

**Supplemental Fig 7: Evaluation of CellFuse predicted cell type for IMC breast cancer data**

**A.** Confusion matrix of CellFuse predicted cell types while using expert annotated cells as ground truth.

**B.** Visual inspection of six patients. For each patient, cells in the left panel are colored by ground truth annotations, and those in the right panel are colored by CellFuse predicted labels
